# Stimulus-repetition effects on macaque V1 and V4 microcircuits explain gamma-synchronization increase

**DOI:** 10.1101/2024.12.06.627165

**Authors:** Christini Katsanevaki, Conrado A. Bosman, Karl J. Friston, Pascal Fries

## Abstract

Under natural conditions, animals repeatedly encounter the same visual scenes, objects or patterns repeatedly. These repetitions constitute statistical regularities, which the brain captures in an internal model through learning. A signature of such learning in primate visual areas V1 and V4 is the gradual strengthening of gamma synchronization. We used a V1-V4 Dynamic Causal Model (DCM) to explain visually induced responses in early and late epochs from a sequence of several hundred grating presentations. The DCM reproduced the empirical increase in local and inter-areal gamma synchronization, revealing specific intrinsic connectivity effects that could explain the phenomenon. In a sensitivity analysis, the isolated modulation of several connection strengths induced increased gamma. Comparison of alternative models showed that empirical gamma increases are better explained by (1) repetition effects in both V1 and V4 intrinsic connectivity (alone or together with extrinsic) than in extrinsic connectivity alone, and (2) repetition effects on V1 and V4 population input rather than output gain. The best input gain model included effects in V1 granular and superficial excitatory populations and in V4 granular and deep excitatory populations. Our findings are consistent with gamma reflecting bottom-up signal precision, which increases with repetition and, therefore, with predictability and learning.

**Highlights:**

- We model learning effects in macaque visual cortex using Dynamic Causal Modeling.
- Microcircuit-level changes explain the repetition-induced gamma increases.
- The best models include changes 1) within V1 and V4 and 2) in neuronal input gain.
- Gamma may reflect bottom-up signal precision.

## 1. Introduction

During natural viewing, animals repeatedly encounter the same scenes, shapes, colors, and patterns. These repetitions constitute statistical regularities in space and time that can be learned by the brain. Learning natural image statistics can serve to make better predictions about the environment (Bastos et al., 2012; Friston and Kiebel, 2009; Rao and Ballard, 1999; Srinivasan et al., 1982) and to encode information about it more efficiently (Ay et al., 2008; Gershman and Niv, 2010; Schwartenbeck et al., 2019; Smith et al., 2020; Tervo et al., 2016; van Leeuwen, 1990). The learning process begins when the animal is presented with an unforeseen stimulus once or a few times (de Gardelle et al., 2013; Desimone, 1996; Friston, 2005). Numerous studies using oddball or roving-standard paradigms (e.g., studying mismatch negativity) have reported transient effects on neuronal responses to the first presentation of a new stimulus (Garrido et al., 2009; Peter et al., 2021; Stauch et al., 2021; Stefanics et al., 2014), which can result from arousal (Stauch et al., 2021) or from rapid changes in synaptic efficacy (e.g., stimulus-specific adaptation). When the efficacy (or precision) of bottom-up prediction-error signals increases relative to those of top-down predictions, the animal can respond more readily to a novel stimulus in the environment.

On a longer timescale, if the stimulus keeps appearing repeatedly, the relative probability of that stimulus and, thereby, the statistics of the environment change. For an animal to behave appropriately within the new environment, sensory learning must take place. This learning requires long-term plastic changes that imprint the content of learned predictions on sensory cortical circuits. In the visual cortex, changes in neuronal responses upon the repetition of visual stimuli have been found at the earliest stages of cortical visual processing in primates, in area V1. Responses in V1 have been shown to be modulated with stimulus repetition in two distinct ways: firing rates decrease (Peter et al., 2021), while simultaneously, gamma synchronization increases (Brunet et al., 2014; Peter et al., 2021) often up to two- or three-fold (**Fig. 1B**). These repetition-induced changes were found using both gratings and naturalistic stimuli (Peter et al., 2021), and the gamma-power increase has also been replicated in a human MEG study using gratings (Stauch et al., 2021). The repetition-related change of gamma is specific for the repeated stimulus and persists over some time, as one would expect for a learning process. Finally, the repetition-related gamma-power increase has also been reported for area V4, along with increased gamma coherence between V1 and V4 (Brunet et al., 2014). Superficially, there seems to be a disparity between the repetition-related changes of gamma power and firing rates. However, it has been proposed that increased synchronization in V1 can compensate for reduced firing rates by rendering spikes more efficient (Gotts, 2003; Gotts et al., 2012). This interpretation is consistent with the increased behavioral performance in tasks with repeated stimuli and with the notion that learning image statistics is essential within the predictive coding framework.

**Figure 1.**
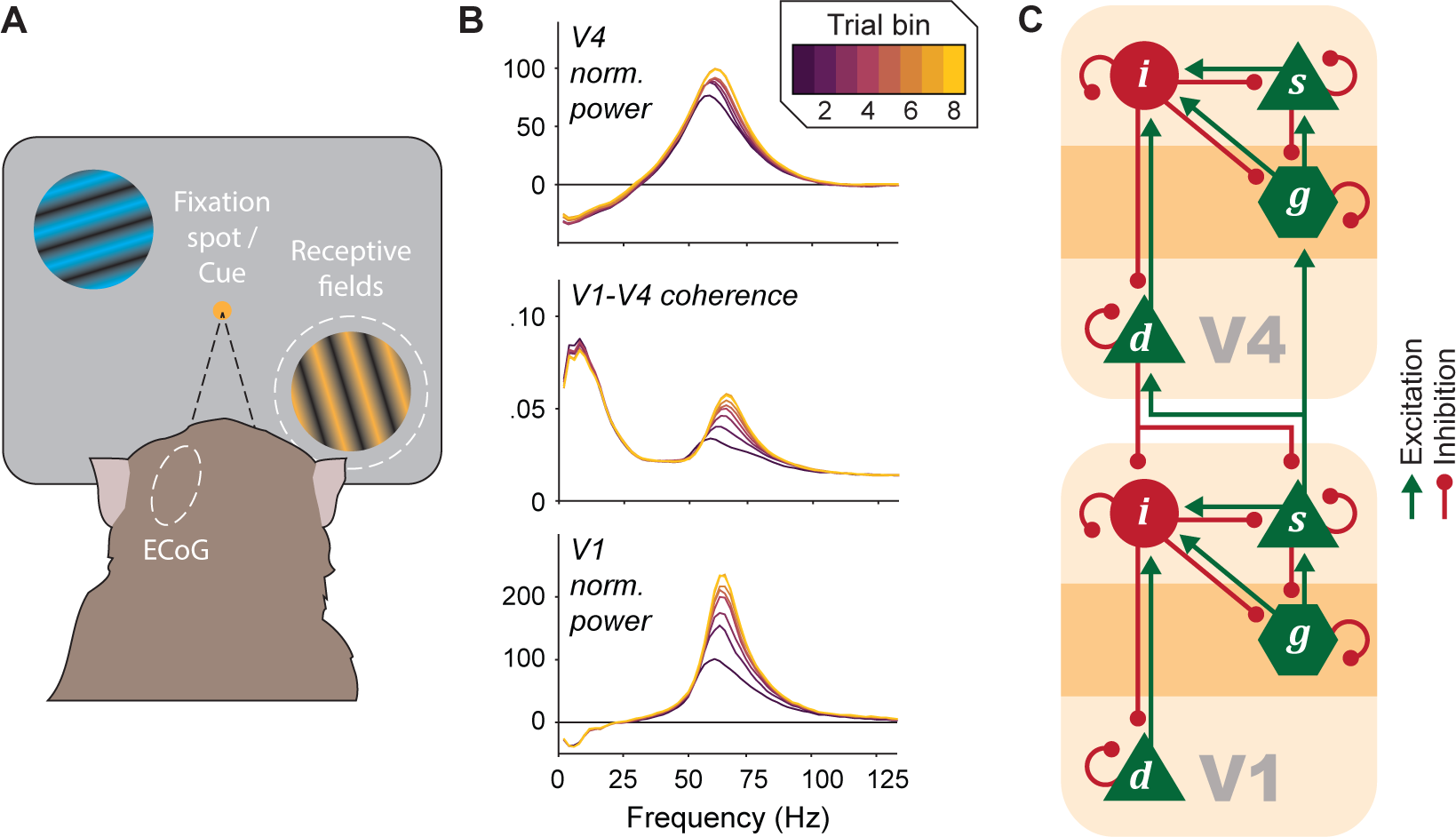
Dynamic causal modelling of repetition-induced changes in the power of V1 and V4 LFP power and V1-V4 coherence spectra. **(A)** The animal performed a selective attention task (Bastos et al., 2015b). In short, the monkey held its gaze at the fixation dot in the center of the screen, while attending to one of two gratings that appeared at equal eccentricities (illustration not to scale). In the current analysis, epochs were pooled over the two attentional conditions, so that the attentional aspect of the task is irrelevant. **(B)** V1 (*bottom*) and V4 (*top*) power change relative to pre-stimulus baseline, and V1-V4 coherence (*middle*), averaged over all visually driven sites or site pairs, respectively, and all sessions of monkey P. The epochs of each session were split into 8 equally sized non-overlapping bins according to their order within the session, and power or coherence spectra were calculated per bin. Modified with permission from Brunet et al. (2014). **(C)** Schematic illustration of the V1-V4 DCM that was inverted to fit the empirical CSDs, showing the pattern of intrinsic (within-area) and extrinsic (between-area) connectivity. The change of the DCM CSDs from early to late epochs is modeled by a vector of repetition effects that multiply the vector of all intrinsic and extrinsic connection strengths. The four extrinsic connections are illustrated by two bifurcating arrows, and on each of the two a single repetition effect is applied.

Here, we focus on the mechanisms underlying the changes in neuronal synchronization induced by many stimulus repetitions. We aim to illuminate the specific microcircuit-level effects in intrinsic (within-area) and extrinsic (between-area) connectivity that can generate the empirical repetition-induced changes and might be the substrate for learning new sensory input statistics. To this end, we modeled empirical V1 and V4 power spectra and V1-V4 coherence spectra using Dynamic Causal Modelling (DCM) for cross-spectral densities (CSD) (Friston et al., 2012). We used the dataset for which the repetition-related gamma increase was originally reported (Brunet et al., 2014). We simulate the repetition-induced changes of V1 and V4 responses by fitting the DCM to the electrophysiological data. We use the ensuing model to identify the loci of synaptic modulation by stimulus repetition within V1, V4, or the extrinsic connectivity between them. Then, we proceed to further characterize the contribution of distinct intrinsic and extrinsic connections to repetition-related changes in the empirical neuronal dynamics and to identify the most concise set of repetition effects that can explain the experimental data. Finally, we discuss our findings in the context of the predictive coding framework, where an increase in gamma synchronization can reflect a learning-induced increase in the precision of bottom-up prediction errors.

## 2. Methods

### 2.1. Experimental details

The animal ethics committee of Radboud University (Nijmegen, the Netherlands) approved all animal experimental procedures. Data from two adult male Rhesus monkeys (*Macaca mulatta*) were used in this study. The monkeys were trained to perform a covert selective attention task (**Fig. 1A** bottom). The attention contrast has not been used in this paper; but task details are given here for completeness. The monkeys had to touch a lever and fixate on a spot that appeared in the center of a screen. At a variable time, one blue and one yellow drifting circular grating appeared at equal eccentricities from the fixation spot. The gratings had a diameter of 3°, the luminance within each grating was defined by a sine-wave function with a spatial frequency of ≈1 cycle/degree and a drift velocity of ≈1 degree/s and the two gratings were physically isoluminant. Also, at a variable time, the fixation spot changed color to either blue or yellow, cueing the grating with the same color to be attended. At a variable time after the cue appeared, the curvature of one of the gratings changed transiently, with an equal probability that the target or the distracter changed. The monkey had to respond to changes in the target by releasing a lever, after which the trial ended, and the monkey obtained a fluid reward.

Electrocorticography (ECoG) grids, with 252 electrode contacts, were implanted in the left hemispheres of the two monkeys. They were used to collect Local Field Potential (LFP) data while the monkeys were performing the attentional task. The signal was low-pass filtered at 250 Hz. In order to remove the recording reference, the signals between neighboring electrodes were subtracted in the time domain. The ensuing bipolar derivatives are hereafter referred to as ‘(recording) sites”. For more details about the recordings and the assignment of sites to cortical areas see (Bastos et al., 2015b; Brunet et al., 2014).

### 2.2. Data preprocessing

Data was segmented into non-overlapping epochs of 500 ms, starting either 500 ms after the cue change, or 300 ms after the distractor-change onset. For each site separately, epochs were demeaned, the standard deviation (SD) across all values of all epochs was calculated and used to normalize by dividing all values by this SD.

In order to characterize the increase in gamma power and coherence with stimulus repetition, we enriched our data features by selecting two groups of epochs from each recording session as follows: (1) We discarded the first 10 epochs from each session, because previous studies (Peter et al., 2021; Stauch et al., 2021) have shown that there can be a rapid initial drop in gamma power, and this was not the phenomenon we wished to characterize. These 10 epochs corresponded to 2-6 trials per session (38 trials in total) in monkey K, and 3-5 trials per session (42 trials in total) in monkey P. (2) From the remaining epochs of a session, we selected the first 100 epochs, referred to as the ‘early’ condition, and the last 100 epochs, referred to as the ‘late’ condition. (3) We separately pooled the early epochs and the late epochs over sessions (monkey P: 11 sessions, 1100 epochs per condition; monkey K: 9 sessions, 900 trials per condition). If a recording session did not contain at least 210 epochs, it was excluded from the analysis (1 session in monkey P).

We pre-whitened the data in order to remove the 1/*f* component and accentuate the gamma component in the power and coherence spectra, as this was the data feature of interest. Before pre-whitening, we downsampled the LFP to 250 Hz. Then we used the ARfit toolbox (Neumaier and Schneider, 2001; Schneider and Neumaier, 2001) to calculate the single coefficient *r* that parameterizes the first-order auto-regressive (AR) model to fit the data. The coefficient was used to calculate the whitened signal samples as:

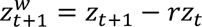

where *z* and *z*^*w*^ are the samples of the original and whitened data respectively and *t* denotes time steps. After pre-whitening, we used the BSMART toolbox (Cui et al., 2008) to fit an 8-order AR model to the data and to estimate the CSD data features that were then fitted by the DCM. In our plots we show the power and coherence from the data-fitted AR model and those predicted by the DCM. Note that all displayed coherence spectra show the coherence in terms of its magnitude, as reported in Brunet et al. (2014) and as opposed to magnitude squared used in some other reports.

### 2.3. V1-V4 site pairs

To study the repetition-related changes on gamma power and coherence, we selected V1-V4 site pairs, which were most visually responsive and showed the strongest coherence. This selection used the pooled data from all trials to avoid selection bias. To quantify how responsive each site was, we calculated the stimulus-induced power change, i.e. the percent change relative to a pre-stimulus baseline (500-0 ms before stimulus onset), and summed those values over a frequency range of 52-74 Hz for monkey P and 68-82 Hz for monkey K (same frequency bands as used in Brunet et al. (2014)) to obtain their ‘visual gamma’. Then, for each monkey, we performed the following selection: (1) We used the visual gamma to select the 20 most driven V1 sites and 5 most driven V4 sites. (2) We formed all possible combinations between them resulting in 100 V1-V4 pairs. (3) We measured—per V1-V4 pair—the V1-V4 gamma coherence within the same frequency range as for the gamma power and selected the 25 pairs with the strongest total gamma coherence. Thus, in total 50 site pairs were selected for modelling.

### 2.4. Dynamic Causal Modelling

We fitted the cross spectral density of the selected V1-V4 pairs with DCM for CSD. DCM for CSD is a well-established model for EEG/MEG and LFP responses, as it describes the mesoscopic scale of neuronal activity and operates on frequency domain data features, making use of both the absolute value and the argument of the complex cross-spectral density matrix (Friston et al., 2012). It can therefore be used to explain neuronal oscillations in terms of their relative amplitude and, crucially, phase, and to infer repetition-induced changes in intrinsic and extrinsic connectivity parameters.

Our DCM comprised two nodes, one for V1 activity and one for V4. Within each node, neuronal populations were modelled using neural masses (David and Friston, 2003; Friston et al., 2012; Jansen and Rit, 1995). The dynamics of each population are determined by two operations: the transformation of average presynaptic firing rates to an average postsynaptic depolarization—by convolution with an alpha function—and the transformation of the postsynaptic depolarization to the firing rate output of the population using a sigmoid function. The two nodes had the same four neuronal populations: an excitatory population in the input (granular) layer (***g***), two pyramidal populations in the superficial (***s***) and deep (***d***) layer and an inhibitory population (***i***) (Auksztulewicz and Friston, 2015; Bastos et al., 2015a; Bastos et al., 2012). The model also described the intrinsic and extrinsic connectivity within and among populations, respectively. This model has been previously validated with regard to the anatomy of the early visual cortex and the functional architectures implied by predictive coding schemes (**Fig. 1C**). The inhibitory connections that stem from excitatory populations can be read as being mediated indirectly by local inhibitory populations (that are not included explicitly in the model for simplicity). This ensures excitation-inhibition balance in the network, and that the microcircuit dynamical system has a fixed-point attractor around which it can be expanded (Bastos et al., 2015a; Moran et al., 2007).

An observation model accounts for the measurement of the data by modeling the LFP as the sum of a neuronal component, which is the weighted sum of the depolarizations of all contributing populations, and channel-specific and non-specific noise that is inherent to the recording. The specific observation model used here included a parameter that accounted for pre-whitening of the data. In this study, only the superficial pyramidal population contributed to the LFP as described previously in Katsanevaki et al. (2023).

Because of the stochastic nature of the dynamical model, we can only identify the model *m* and parameters *θ* that are *more likely* to give rise to the empirical CSD data *y*, i.e., for which there is more evidence based on the data. The calculation of both the evidence *p*(*y*|*m*) and of the posterior distribution *p*(*θ*|*y*, *m*) requires a specification of the joint probability *p*(*y*, *θ*|*m*), known as the generative model:

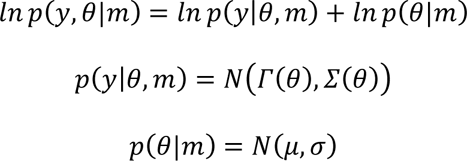

The first term on the right side of the first equation is the likelihood of the data given the model and the parameters, and the second term is the prior distribution over the parameters. The likelihood is defined as a multivariate normal distribution *N* with mean *Γ*(*θ*) and covariance matrix *Σ*(*θ*). *Γ* denotes the nonlinear model that generates the CSD data and is parameterized by *θ*, i.e., the set of differential equations that comprise the neuronal and observation model. The mean and variance of the observation model act as hyperparameters. The prior mean and variance of all parameters *θ* (including the hyperparameters) are given in **Table 1**.

**Table 1.**
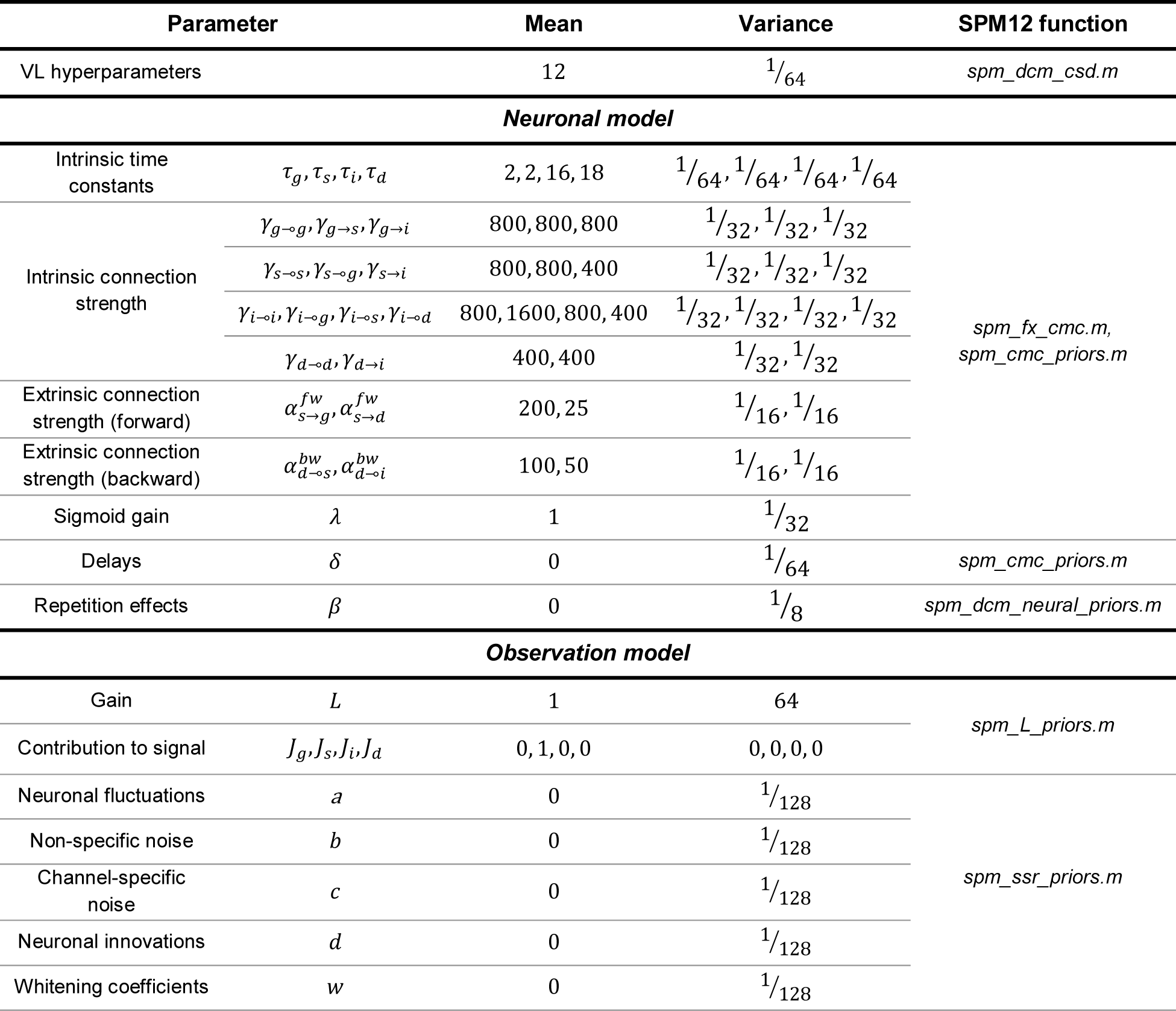
Prior distribution mean and variance of all DCM model parameters.

To obtain an approximation *q*(*θ*) of the true posterior over parameters *p*(*θ*|*y*, *m*) one can perform gradient ascent on the variational negative free energy *F*. *F* is a logarithmic quantity that is equal to the log evidence of the data minus the gap (in the form of a Kullback-Leibler divergence) between the approximate and the true posterior parameter distribution (*q*(*θ*) and *p*(*θ*|*y*) respectively):

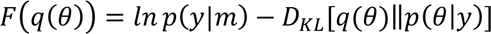

Conditioning the terms on the model has been omitted for simplicity. Since the evidence is fixed but unknown, maximizing *F* minimizes the divergence term, i.e., improves the approximation to the posterior over parameters, and *F* better approximates the log model evidence. *F* is maximized using the standard Variational Laplace (VL) scheme described in detail elsewhere (Friston et al., 2007; Zeidman et al., 2023). In brief, VL performs gradient ascent in alternating steps of optimizing the parameters to maximize *F*, while fixing the hyperparameter estimates, and then optimizing the hyperparameters to maximize *F*, while fixing the parameter estimates.

Because *F* is a good approximation to the log model evidence, it can be used to score different models by how well they explain the data. This is easier to understand if the above definition of *F*is rearranged as:

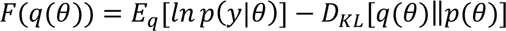

This expression shows that *F* is a trade-off between the model *accuracy* (expected log likelihood of the data) and the model *complexity* (the divergence of the posterior parameter estimates from their prior densities).

As in our previous study (Katsanevaki et al., 2023), we used a mean and variance for the hyperparameters that reflect both the high precision (low variance) of the ECoG measurement—relative to EEG or MEG data—and the averaging implicit in computing the complex cross spectra. The data features on which the DCMs were inverted were a weighted mixture of (1) the V1-V4 complex CSD values, (2) the cross-covariance function and (3) the MVAR coefficients estimated from the data, weighted with relative precisions of 1, 1 and 1/8, respectively. We specified a window of 0-100 Hz as the frequencies of interest for inverting the DCM. We used MATLAB and the SPM12 toolbox for specifying and inverting the DCMs and subsequent analyses. The SPM12 functions containing all model parameters and specifications are mentioned in **Table 1**.

### 2.5. Quantification of repetition effects

Within the DCM we have defined, changes between the early epochs and the late epochs were modelled by a vector of repetition effects (*β*) that modulate the strength (*γ*) of the 24 intrinsic and 4 extrinsic connections. The two forward and backward connections were modulated by a single *β* value each, giving a total of 26 *β* values.

Generally, to ensure that scale parameters (e.g., rate and time constants) have non-negative values, all parameters (except the electrode gains) have log-normal distributions and log-normal priors. This means that connection strengths in the early epochs were specified as *e*^*γ*^, where *γ* is the parameter that is being estimated when a DCM is inverted. The repetition effects are added to the early-epoch connection strengths, resulting in connection strengths *e*^*γ*+*β*^ in the late epochs. The relative ratio of ‘late’ to ‘early’ connection strength is:

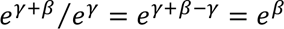

Thus, positive and negative *β* values mean that parameters increase or decrease with repetition, respectively.

To combine results over the 50 fitted V1-V4 pairs, we used Parametric Empirical Bayes (PEB) (Friston et al., 2016; Zeidman et al., 2019), considering each pair as a ‘unit’. PEB assumes that all units have the same neuronal architecture but differ in their specific parameter values, e.g., connection strengths or condition-specific changes. This means that one can build a hierarchical model with multiple levels, where the parameter values of each level are encoded by a Gaussian distribution whose mean and variance are generated by the level above. These constitute empirical priors for the DCM parameters, which effectively shrink the individual unit estimates around a group mean, in proportion to the confidence about the unit estimates. Crucially, PEB takes the posterior covariance (and not just the posterior means) of parameters to the group level, which means that units with more precise estimates will contribute more to the group mean (Friston et al., 2016; Zeidman et al., 2019).

In our case, the parameters of interest are the repetition effects *β*, for which PEB is defined as:

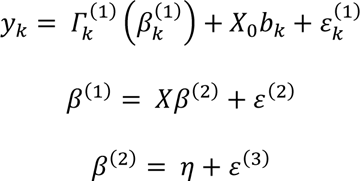

The superscripts in parentheses indicate the PEB level. The first row represents the first hierarchical level of PEB, and it simply describes how the data *y*_*k*_ of each unit *k* is generated: *Γ_k_*^(1)^ is the nonlinear model, i.e. DCM, of that unit with parameters *β_k_*^(1)^ that generates the data, parameters b*_k_* and design matrix *X*_0_ specify a general linear model (GLM) that models uninteresting effects, such as the signal mean, and *ε_k_*^(1)^is the unit-specific zero-mean additive random or sample effect (c.f., observation or sample noise). The second and third row describe the level above: the parameters *β*^(1)^ that specify the first-level models are generated by a GLM with group-level parameters *β*^(2)^, design matrix *X* and between-unit variability *ε*^(2)^. The second-level parameters are sampled from a distribution with mean *η* and variability *ε*^(3)^, as shown in the last equation. In effect, Parametric Empirical Bayes is the (Bayesian) generalization of general linear models of random effects.

### 2.6. Sensitivity analysis

To assess the contribution of specific model parameters (e.g., connectivity) to measurable responses (i.e., gamma activity), we perturbed the parameter estimates—from the full DCM—by adding a value ranging from -0.1 to +0.1 in steps of 0.025 to the posterior expectation of individual parameters expressing repetition effects. We then calculated the new cross-spectral densities of the modified DCM and plotted them for each perturbation. We show the sensitivity to the perturbation of all parameters that showed significant repetition effects following PEB. We did this analysis for two representative DCMs, one from each monkey, which were closest to the PEB estimates for the repetition effects. For this, we calculated for each of the 50 DCMs the total absolute difference between all repetition effects in the DCM and in the PEB and selected for the sensitivity analysis, per monkey, the DCM with the lowest total difference (these DCMs from the two monkeys also happened to be the two DCMs with the overall lowest total difference when considering both animals together).

This sensitivity analysis allows one to see how measurable DCM features, i.e., cross spectral densities, are caused by underlying changes in key model parameters, i.e., connectivity. The trends shown in Fig. 3 and Fig. S3 are not necessarily the same in the rest of the DCMs. Also, they would not necessarily be the same if all or some other repetition-related (or any other) parameters were perturbed simultaneously. This analysis only aims to quantify how individual parameters shape the power- and cross-spectra.

**Figure 2.**
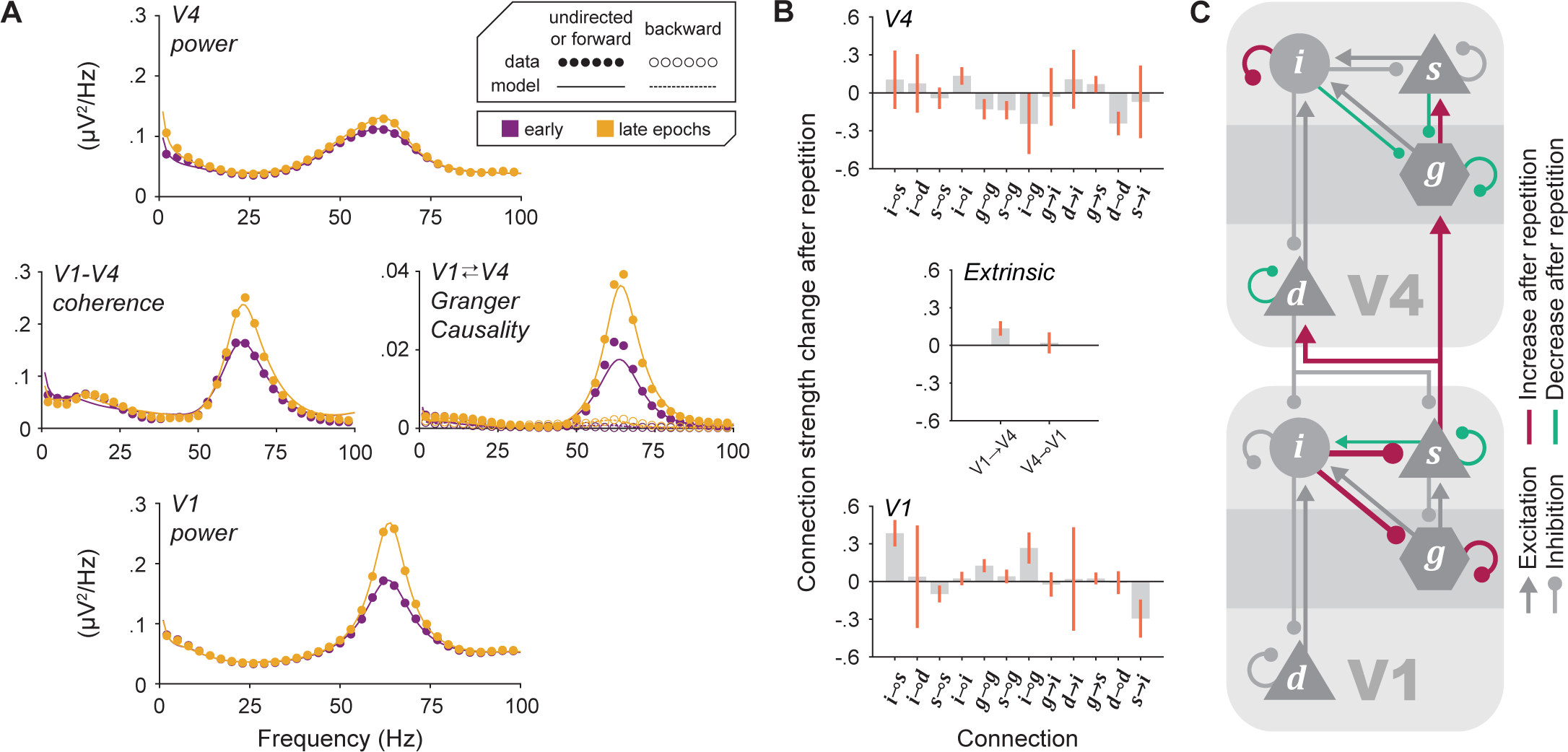
The V1-V4 DCM can reproduce the data features. **(A)** Empirical (*circles*) and predicted (*lines*) spectra of V1 power (*bottom*), V1-V4 coherence and GC (*middle*) and V4 power (*top*), averaged over all V1-V4 site pairs and DCMs of monkey P. The empirical CSD spectra were averaged over 100 early epochs (*purple*; first epochs in each recording session after discarding the first 10 epochs) and 100 late epochs (*yellow*; last epochs in each session), respectively. **(B)** Group-level PEB posterior logarithmic estimates and 95% credible intervals (*orange lines*) of the repetition effects on intrinsic V1 (*bottom*), extrinsic (*middle*) and intrinsic V4 (*top*) connection strengths. If a credible interval does not cross the zero line, the effect is significant in the sense that there is strong evidence for it (Kass and Raftery, 1995). **(C)** Schematic illustration of the significant positive (*magenta*) and negative (*teal*) repetition effects from panel (B). The ratio of each colored line width to the width of the grey lines is equal to the ratio of ‘late’ to ‘early’ connection strength for that connection.

**Figure 3.**
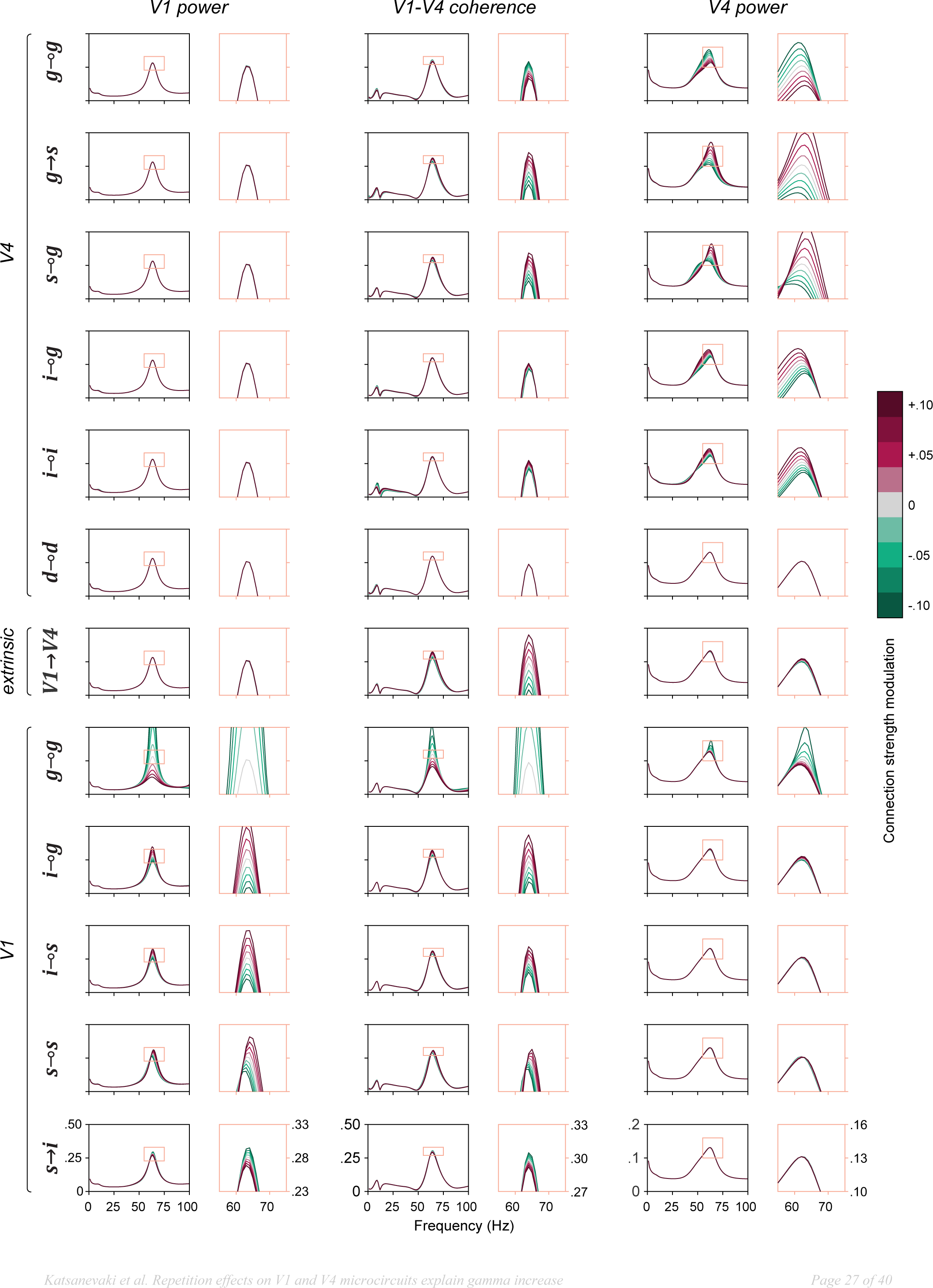
Sensitivity of V1 power (*left*), coherence (*middle*) and V4 power (*right*) spectra to small perturbations of the strength of individual connections in a representative DCM of monkey P. Grey lines correspond to the single DCM spectra in the ‘late’ condition; there is no perturbation of the connection strength apart from the repetition effect. Each row shows the change of the power and coherence spectra induced by the perturbation of an individual connection strength (connection label on the left side). The perturbation consists of adding a constant ranging from −0.1 (*teal*) to +0.1 (*magenta*) in steps of 0.025 to the repetition effect of the connection (the total value is applied as an exponent to a multiplier of the connection strength). Note that this sensitivity analysis generates combinations of parameters that are not constrained by the data spectra.

### 2.7. Bayesian Model Reduction (BMR)

BMR allows to infer the posterior parameter distributions and evidence for a DCM from the posteriors and evidence of a previously inverted DCM, provided the two models (the new DCM and the inverted DCM) differ only in their priors. For example, a reduced DCM with fewer parameters (where some parameters have zero mean and variance) or more restrictive priors (where some parameters have lower variance) can be easily computed from a more expressive model (in our case, this is the full model). With this in mind, BMR can be used to estimate the contribution of specific parameters to model evidence.

In our case, any DCM in which only a subset of intrinsic or extrinsic connections can be modulated by stimulus repetition is a reduced model relative to the full model. For each hypothesis we wanted to test, we removed relevant repetition effect combinations from the model, e.g., extrinsic and intrinsic V4 repetition effects. Then, we compared the evidence of all ensuing reduced models, the full model, and a null model without any repetition effects. This enabled us to assess the relative evidence for different sets of parameters in explaining repetition-induced changes in the empirical spectra.

### 2.8. Comparing DCMs with Bayes factors

Bayes factors can be used to assess the relative evidence for alternative DCMs in an intuitive manner. The Bayes factor in favor of one model against another is the ratio of their model evidences, i.e., the probability to observe data *y* given each model. Since *F* is the approximation of evidence, the logarithmic Bayes factor in favor of model *i* against model *j* is simply the *F* difference between two models:

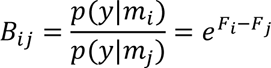

One can calculate the posterior probability of each model using *F* (or *B*). Assuming all models have equal prior probabilities, the posterior probability of model *i* is:

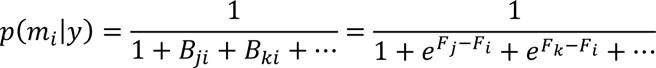

where *i*, *j*, *k*, … are all models that can explain the data *y*. Under a uniform prior over models, the Bayes factor is also equivalent to the ratio of posterior model probabilities. *B*_*ij*_ = 20 means that model *i* is 20 times more probable than model *j* given the data, or in other words, a 95% belief that model *i* is true. This corresponds to free energy difference *F*_*i*_ − *F*_*j*_ = *lnB*_*ij*_ ≈ 3. For each comparison, we report *lnB*^−1^, the logarithmic Bayes factor in favor of each model against the best model (that has the maximum posterior probability), which is more negative for less probable models. We use the Bayes factor interpretations of (Kass and Raftery, 1995) but provide a slightly modified version to show *lnB*^−1^for ease of reference (**Table 2**).

**Table 2.** Interpretation of Bayes factors. Slightly modified from Kass and Raftery (1995) to show *lnB*^−1^values instead of corresponding 2*lnB* values. Here, *B* is the Bayes factor in favor of the *m*_*MP*_ (model with the highest posterior probability) in a given comparison against another model *m*, which can be interpreted as a posterior odds ratio or ‘how many times more likely is the *m*_*MP*_ compared to model *m*’. Therefore, the last column also refers to the evidence in favor of the *m*_*MP*_ against model *m*. Finally, *lnB*^−1^ is essentially the free energy difference between the *m*_*MP*_ and another model *m*.

| $B$ | $\ln B^{-1}$ | Evidence against $m$ |
| --- | --- | --- |
| 1 to 3 | -1 to 0 | Not worth more than a bare mention |
| 3 to 20 | -3 to -1 | Substantial |
| 20 to 150 | -5 to -3 | Strong |
| > 150 | < -5 | Very strong |

### 2.9. Bayesian Model Averaging (BMA)

To estimate the magnitude of interesting parameters, i.e., repetition-induced changes, when the evidence of any single model is not definitive, it can be useful to average the posterior estimates of the parameters over multiple models. This can be done with BMA, where each posterior parameter estimate is weighted by the posterior probability (marginal likelihood) of the respective model (Madigan and Raftery, 1994):

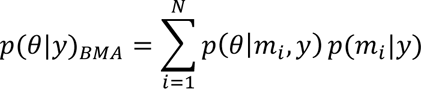

It is considered that it is most meaningful to calculate the BMA over models whose posterior probability relative to the best model (with the maximum posterior probability, *m*_*MP*_) is within a predefined Occam’s window (Madigan and Raftery, 1994; Penny et al., 2010). Here, we have set the limit for Occam’s window to *lnB*^−1^ ≥ −5 so that we consider all models against which the evidence is less than ‘very strong’ compared to the *m*_*MP*_ model. The results were virtually the same when a stricter window of *lnB*^−1^ ≥ −3 was used.

## 3. Results

We modelled electrophysiological data from two monkeys presented with repeating stimuli (**Fig. 1A**, section 2.1). It has been previously shown that the repetition of stimuli within a recording session results in an increase of V1 and V4 LFP gamma power and an increase of V1-V4 LFP gamma coherence (**Fig. 1B**) (Brunet et al., 2014). In the current study, we characterize the mechanisms underlying the repetition-induced changes in the V1 and V4 spectra by fitting a V1-V4 DCM (**Fig. 1C**) to the empirical CSD. After discarding the first 10 epochs of each session (see section 2.2), the CSD was averaged over the following 100 epochs, referred to as the ‘early’ condition, and over the last 100 epochs, referred to as the ‘late’ condition. The difference between the average CSD from the two conditions exhibited the previously reported repetition-increase in local and inter-areal gamma synchronization.

### 3.1. DCM reproduces the empirical repetition-induced changes

For each of the two monkeys, 25 pairs of simultaneously recorded V1 and V4 sites were selected, resulting in a total of 50 V1-V4 pairs (see section 2.3). For each pair, the V1-V4 CSD matrix was fitted with a DCM that consisted of a V1 and a V4 node (see section 2.4 and Table 1 for all prior parameter distributions and Fig. S1 for 5 randomly selected example pairs per animal). Power and coherence were averaged over the 25 V1-V4 pairs and 25 V1-V4 DCM-predicted spectra from each monkey separately to show how closely the predicted spectra fit the observed empirical spectra and that they replicate the effects of stimulus repetition (**Fig. 2A, S2**). The full spectra were not averaged over the two monkeys, because they exhibited different gamma peak frequencies.

DCM allows one to quantify directed functional connectivity between V1 and V4 in the form of Geweke’s Granger Causality (GC) (Geweke, 1982). GC measures the reduction of unexplained variance in the AR model of the receiver, when it is computed using information from the sender and the receiver, as opposed to when it is computed from using information from the receiver alone (Bressler and Seth, 2011; Chicharro, 2011). The V1-to-V4 GC estimated directly from the empirical data exhibited a clear peak in the gamma-frequency range, both for the early and the late epochs. The comparison between the conditions revealed that stimulus repetition resulted in GC increases, which also peaked in the gamma-frequency range. The GC derived from the DCM closely reproduced all these features. The V4-to-V1 gamma GC was much weaker than the V1-to-V4 gamma GC (Bastos et al., 2015b; Michalareas et al., 2016; van Kerkoerle et al., 2014) but showed qualitatively similar changes from early to late epochs, which were also well explained by DCM.

By fitting 50 DCMs, we obtained 50 sets of posteriors that can generate the empirically observed repetition effects of gamma power and coherence. We then used PEB to fit a hierarchical model that could explain the distribution over the posterior parameters of the multiple DCMs (see section 2.5). In this way, we obtained a group-level estimate of the mean and variance of repetition effects on the strength of each intrinsic and extrinsic connection in the V1-V4 microcircuit (**Fig. 2B, 2C**). We found significant repetition effects in V1 (*β*_*i*⊸*s*_ = 0.38 ± 0.06; *β*_*s*⊸*s*_ = −0.1 ± 0.04; *β*_*g*⊸*g*_ = 0.13 ± 0.03; *β*_*i*⊸*g*_ = 0.27 ± 0.07; *β*_*s*→*i*_ = −0.29 ± 0.09), in V4 (*β*_*i*⊸*i*_ = 0.13 ± 0.04; *β*_*g*⊸*g*_ = −0.13 ± 0.05; *β*_*s*⊸*g*_ = −0.14 ± 0.04; *β*_*i*⊸*g*_ = −0.24 ± 0.14; *β*_*g*→*s*_ = 0.07 ± 0.04; *β*_*d*⊸*d*_ = −0.24 ± 0.06), and in extrinsic connectivity (*β*^*fw*^ = 0.14 ± 0.04). These values correspond to the logarithmic ratio (and its variance) of the late-epochs connection strength over the early-epochs connection strength, which means that positive and negative values correspond to increases and decreases with stimulus repetition, respectively.

In order to better understand how the individual repetition-related changes revealed in the PEB model can impact the microcircuit dynamics and especially gamma power and coherence, we performed a sensitivity analysis (see section 2.6). The sensitivity of V1 and V4 power and V1-V4 coherence is shown for two representative DCMs, one from monkey P (**Fig. 3**) and one from monkey K (**Fig. S3**). We observed the following trends:

- Increasing ***s*** self-inhibition in V1 increased V1 gamma power and V1-V4 gamma coherence in monkey P, whereas it decreased V1 gamma power in monkey K.
- Increasing ***s***-to-***i*** excitation in V1 decreased V1 gamma power and slightly decreased V1-V4 gamma coherence in monkey P, whereas it had no observable effect in monkey K.
- Increasing ***i***-to-***s*** inhibition in V1 increased V1 gamma power in both monkeys, and in monkey P increased V1-V4 gamma coherence.
- Increasing extrinsic excitation increased V1-V4 coherence in both monkeys.
- Increasing ***i*** self-inhibition in V4 increased V4 gamma power and V1-V4 gamma coherence in both monkeys.
- Increasing ***d*** self-inhibition in V4 did not have any observable effect on gamma power and might have slightly decreased V1-V4 gamma coherence.
- Increasing ***g*** self-inhibition in V1 or V4 reduced local gamma power and V1-V4 gamma coherence in both monkeys. Increasing the same connection in V1 of monkey P additionally reduced V4 gamma power.
- Increasing other inhibition to ***g*** (***s***-to-***g*** in V4 and ***i***-to-***g*** in V1 and V4) increased local gamma power and V1-V4 gamma coherence in both monkeys.

### 3.2. Extrinsic vs. intrinsic repetition effects

Using the estimated PEB model as a full model, we used BMR (see section 2.7) to remove potentially redundant parameters in order to disclose the most likely concise subset of repetition effects in the microcircuit. BMR is an analytical method to calculate the free energy and posterior estimates for a reduced DCM directly from the free energy and posteriors of the full model, offering an efficient way to test multiple hypotheses about which subset of connection strength changes can best explain the empirical repetition-induced changes.

The first question we asked was whether the extrinsic, the intrinsic V1 and the intrinsic V4 parameter changes all contribute significantly to the repetition effects. To this end, we grouped connections as V1-intrinsic, V4-intrinsic, or extrinsic. Then, we formed all combinations of ‘switching on’ or ‘off’ repetition effects on all connection strengths within each of the three groups (2^3^ = 8 combinations). Finally, we estimated the relative evidence of all models that corresponded to these combinations (see section 2.8) to find the model with the highest posterior probability in light of the data *y* (namely, the marginal likelihood of the model in question, under a uniform prior over all possible models) (**Fig. 4A**). For a given set of models that are being compared, the best model is referred to as *m*_*MP*_. For the comparison presented in this paragraph, the *m*_*MP*_ is the PEB model that allows the full range of repetition effects in V1, V4 and in the extrinsic connections (*p*(*m*_*full*_|*y*) = 0.74). The next best model is the model with only intrinsic repetition effects (*p*(*m*_3_|*y*) = 0.26). These are the only two models with non-trivial posterior probabilities. The evidence in favor of the *m*_*MP*_against the second-best model was, following the definitions in (Kass and Raftery, 1995), ‘not worth more than a bare mention’ (*lnB*^−1^ > −1) (**Fig. 4B**, **Table 3**).

**Figure 4.**
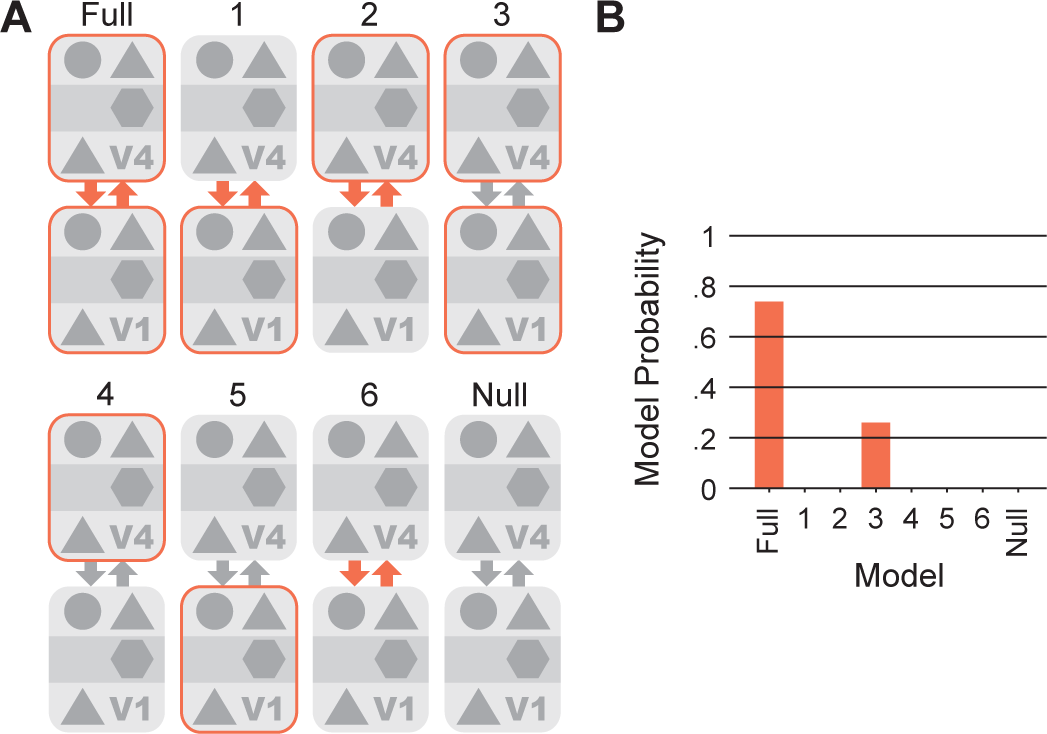
Comparison between the 8 alternative models that allow any combination of the 3 sets of V1 intrinsic, extrinsic and V4 intrinsic repetition effects. **(A)** Schematic illustration of the full, reduced and null models. **(B)** Posterior model probabilities for all models shown in (A).

**Table 3.** Posterior model probabilities and Bayes Factors in favor of each model against the *m*_*MP*_ for the two models with *p*(*m*|*y*) > 0.005.

| Model | $p(m\vert y)$ | $\ln B^{-1}$ |
| --- | --- | --- |
| V1-ext-V4 (full) | 0.74 | 0 |
| V1-V4 (3) | 0.26 | -0.45 |

### 3.3. Repetition effects on neuronal populations

Next, we examined how different populations within V1 and V4 were affected by stimulus repetition. We wished to test two different types of population-level repetition effects, namely the repetition dependent changes in the population input or output gain. To test all possible input-gain models, we grouped intrinsic and extrinsic connections according to which of the eight (V1 or V4) populations they *target* (**Fig. 5A** top), resulting in eight non-overlapping connection groups. We then formed all possible combinations of ‘switching on’ or ‘off’ all repetition effects within each group (2^8^ = 256 combinations). For the output-gain models, we grouped all connections according to which population they *originate from*, and then followed the same logic (**Fig. 5A** bottom). This yielded a total of *N* = 510 reduced models, as the full model and the null model for input gain and the output gain are the same. We compared all models based on their posterior probability, which we report together with the respective logarithmic Bayes factors (against the best model) of all models with a posterior probability higher than 0.005 in **Table 4**. Finally, we did the same analysis but without including extrinsic connections in any of the input or output gain models (*N* = 508) to verify that we get the same results under the second-best model of the previous section, i.e. the model that allowed repetition effects only on intrinsic connections, since the evidence against it was not substantial.

**Figure 5.**
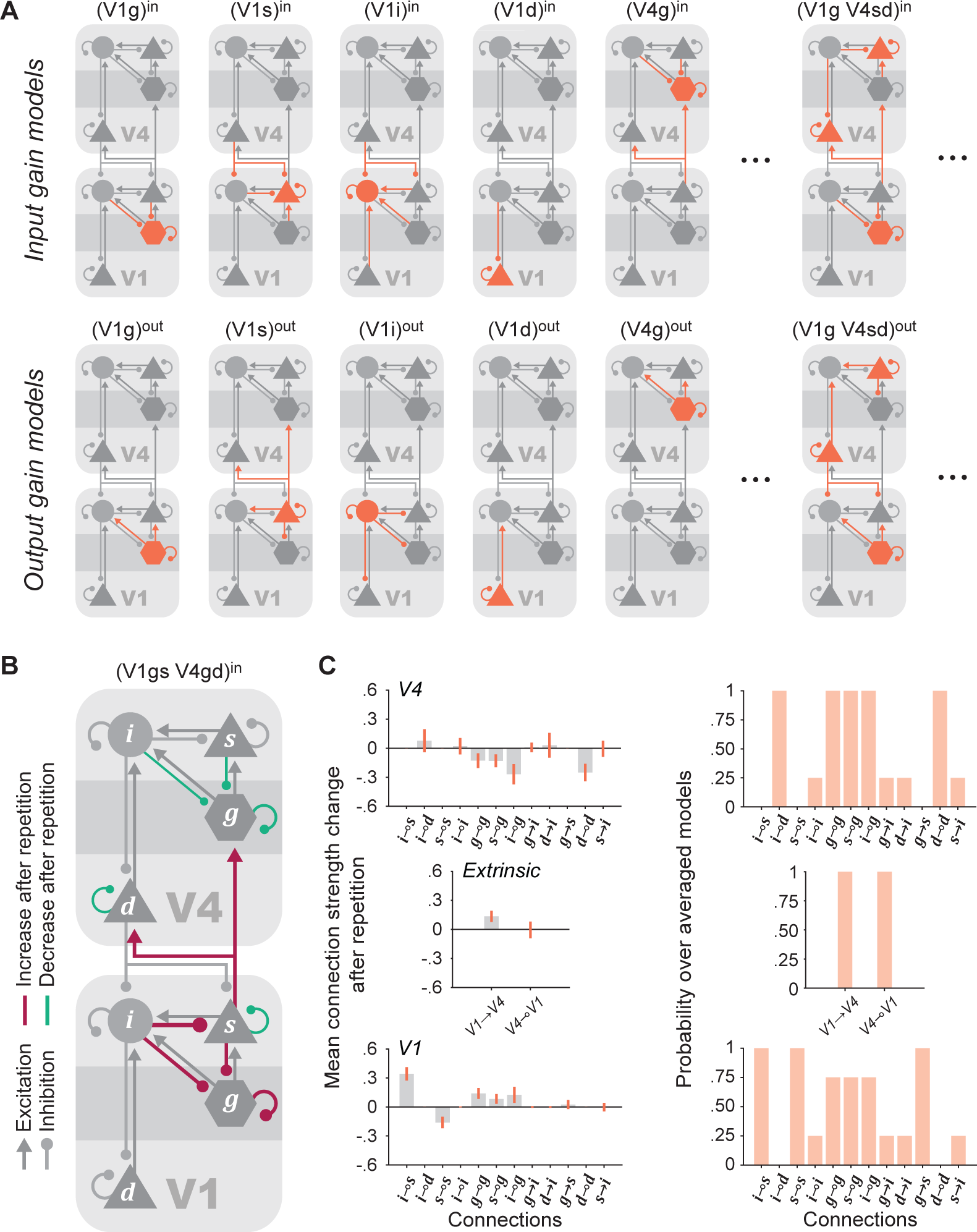
Comparison between all alternative models of population input and output gain in V1 and/or V4. **(A)** Illustration of some input-gain (*top*) and output-gain (*bottom*) models. To test whether repetition modulated the input gain of a population, repetition effects of all afferent connections to that population were allowed. We tested all combinations of the 8 populations. Similarly, to test whether repetition modulated the output gain of a population, we allowed repetition effects in all efferent connections of the population. The comparison was between all input-gain models, all output-gain models, a full model where repetition effects were allowed in all connections, and a null model without any repetition effects. **(B)** Schematic illustration of model (V1gs V4gd)^in^, which was the *m*_*MP*_ amongst all input and output gain models. Colors signify connections with significant positive (*magenta*) and negative (*teal*) repetition effects. The ratio of each colored line width to the width of the grey lines is equal to the ratio of ‘late’ to ‘early’ connection strength for that connection. **(C)** BMA (*left*) and probability (*right*) of repetition effects over all models within an Occam’s window of 5.

**Table 4.** Posterior model probabilities and logarithmic Bayes Factors in favor of each model against the *m*_*MP*_, for the best models of population input or output gain, when extrinsic connections are included (*m*) or excluded (*m*_*ne*_) from the models. All models with *p*(*m*|*y*), *p*(*m*_*ne*_|*y*) > 0.005 are shown in descending probability order.

| Model | $p(m\vert y)$ | $\ln B^{-1}$ | $p(m_{ne}\vert y)$ | $\ln B_{ne}^{-1}$ |
| --- | --- | --- | --- | --- |
| (V1gs V4gd)<sup>in</sup> | 0.810 | 0.00 | 0.826 | 0.00 |
| (V1gs V4gid)<sup>in</sup> | 0.152 | -1.68 | 0.135 | -1.81 |
| (V1gsi V4gd)<sup>in</sup> | 0.008 | -4.56 | 0.008 | -4.60 |
| (V1s V4gd)<sup>in</sup> | 0.006 | -4.96 | 0.008 | -4.67 |
| (V1gs V4g)<sup>in</sup> | 0.005 | -5.07 | 0.005 | -5.03 |

Interestingly, all models with a non-trivial posterior probability (*p*(*m*|*y*) ≥ 0.005) were input gain models. The *m*_*MP*_ for this comparison was (V1gs V4gd)^in^ with *p*(*m*|*y*) = 0.81, which allowed repetition effects in the inputs of the ***g*** and ***s*** populations in V1 and the ***g*** and ***d*** populations in V4 (**Fig. 5B**). This confirms the finding of the previous section, that both V1 and V4 modulations are crucial to explain (or generate) the repetition-induced changes in the empirical spectra. Following the definitions in (Kass and Raftery, 1995), the evidence in favor of the *m*_*MP*_ compared to the next best models is ‘substantial’ against the (V1gs V4gid)^in^ model (−3 < *lnB*^−1^ < −1), ‘strong’ against the (V1gsi V4gd)^in^ and (V1s V4gd)^in^ models (−5 < *lnB*^−1^ < −3) and ‘very strong’ against the (V1gs V4g)^in^ model (*lnB*^−1^ < −5) (**Table 4**, rows 2-3).

We repeated this comparison for models that excluded all extrinsic connections (denoted with the subscript ‘ne’). Here, the *m*_*MP*_was again (V1gs V4gd)^in^ with (*p*(*m*_*ne*_|*y*) = 0.83). The other models with non-trivial posterior probabilities were the same as in the last paragraph, and they had the same evidence relative to the *m*_*MP*_ of this comparison (**Table 4**, rows 4-5).

Comparing the Bayes factors for each model in the case of included and excluded extrinsic connections (**Table 4**, columns 3 and 5 respectively) allowed some additional observations. When the models included repetition effects in the extrinsic connections, we observed that, compared to the *m*_*MP*_: (1) the (V1gs V4gid)^in^ and (V1gsi V4gd)^in^ models have a slightly less negative *lnB*^−1^, and (2) the (V1s V4gd)^in^ and (V1gs V4g)^in^ models have a slightly more negative *lnB*^−1^.

Furthermore, having the posterior parameters of all reduced models, we calculated the BMA of each repetition effect over all models within an Occam’s window (Madigan and Raftery, 1994; Penny et al., 2010) of *lnB*^−1^ > −5. There were *N* = 4 models within Occam’s window for both cases, i.e., including (BMA) or excluding (BMA_ne_) extrinsic connections (**Table 4**, **Fig. 5C**). Finally, we calculated the probability that a given repetition effect was included in those models (**Fig. 5D**). The following repetition effects were significant in the BMA (numbers in parentheses list mean, variance and probability of the BMA_ne_ if they differed from BMA) in V1: *β*_*i*⊸*s*_ = 0.34 ± 0.04 with *p*(*β*_*i*⊸*s*_|*y*) = 1, *β*_*s*⊸*s*_ = −0.16 ± 0.04 with *p*(*β*_*s*⊸*s*_|*y*) = 1, *β*_*g*⊸*g*_ = 0.14 ± 0.03 with *p*(*β*_*g*⊸*g*_|*y*) = 0.75, *β*_*s*⊸*g*_ = 0.09 ± 0.03 with *p*(*β*_*s*⊸*g*_|*y*) = 0.75 (0.08 ± 0.03, 0.75), *β*_*i*⊸*g*_ = 0.12 ± 0.05 with *p*(*β*_*i*⊸*g*_|*y*) = 0.75; in extrinsic connectivity: *β*^*fw*^ = 0.13 ± 0.04 with *p*(*β*^*fw*^|*y*) = 1 and in V4: *β*_*g*⊸*g*_ = −0.13 ± 0.05 with *p*(*β*_*g*⊸*g*_|*y*) = 1, *β*_*s*⊸*g*_ = −0.13 ± 0.04 with *p*(*β*_*s*⊸*g*_|*y*) = 1, *β*_*i*⊸*g*_ = −0.27 ± 0.07 with *p*(*β*_*i*⊸*g*_|*y*) = 1 (−27 ± 0.06, 1), *β*_*d*⊸*d*_ = −0.25 ± 0.05 with *p*(*β*_*d*⊸*d*_|*y*) = 1 (−0.25 ± 0.06, 1).

## 4. Discussion

In summary, we used DCM for CSD to model power and coherence spectra obtained from 50 V1-V4 pairs, 25 from each of two awake macaque monkeys presented with moving grating stimuli for several hundred repetitions. These stimulus repetitions led to increasing gamma power and coherence, as reported previously (Brunet et al., 2014) and replicated in subsequent studies (Peter et al., 2021; Stauch et al., 2021), yet the underlying mechanisms were so far unclear.

The DCM was able to fit the empirical spectra well and to replicate repetition-induced changes in the V1 and V4 dynamics. That is, the model CSD had a spectral profile very close to the empirical CSD, showing clear gamma peaks in V1 and V4 power spectra and in V1-V4 coherence and GC spectra, and these gamma peaks showed increases from early to late epochs. The PEB estimate of the repetition effects over all DCMs included significant effects in intrinsic V1 and V4 connection strengths and a significant increase in the forward connection from V1 to V4. To further characterize the possible contributions of the different connection strength changes to the empirical repetition-induced changes, we performed a sensitivity analysis. This exhibited how the perturbation of individual connections can affect the power and coherence spectra of representative DCMs (we discuss the results in section 4.2).

Then, we computed the model evidence for alternative models that could explain the data using BMR and compared them based on their posterior probabilities. The first test revealed that the repetition-related changes in the power and coherence spectra could be explained best by the models that included repetition effects in the strength of both V1 and V4 intrinsic connections, either in isolation or together with the extrinsic connections between V1 and V4. The second test examined which of all possible reduced models of repetition effects on the input or output gain of any subset of neuronal populations could best explain the repetition-related changes in the spectra. This was done while either including or excluding extrinsic connections, and in both cases, all models with *p*(*y*|*m*) > 0.005 were input gain models. The model with the highest posterior probability included repetition effects on the input gain of the V1 ***g*** and ***s*** populations and the V4 ***g*** and ***d*** populations. Some of the other models in Occam’s window included repetition effects in the ***i*** population in V4 or in V1 or excluded effects in the V1 ***g*** or V4 ***d*** population.

### 4.1 Technical considerations

As is true for all models, our results are contingent on our choices of model and assumptions, as we have discussed previously (Katsanevaki et al., 2023). These choices include the following: (1) the canonical microcircuit model we employed strikes a balance between a simple and a biologically plausible cortical microcircuit, which we deemed appropriate for the given ECoG data; (2) neural mass models apply to the level of neuronal populations, not individual neurons; (3) the model fitting aims at finding the simplest explanation for both the spectra and their repetition-related modulation, which is not necessarily the same as the true (unknowable) explanation.

We found different effects on the intrinsic connectivity in V1 versus V4, both qualitatively (the direction of changes) and quantitatively. These differences might be due to differences in the empirical data from V1 and V4, and/or to the fact that the extrinsic connectivity between V1 and V4 was asymmetric by design. This could play a role in the balance of positive and negative effects between intrinsic and extrinsic connections. It will be a fruitful avenue for future research to include data from areas higher than V4 and extend the model accordingly.

### 4.2 Making sense of sensitivity (in light of PEB and BMR)

In this section, we discuss the insights that can be gained from the results of the sensitivity analysis. The starting point of this study was the increase in local power and inter-areal coherence induced by stimulus repetition, the underlying mechanisms of which we wished to understand (the previously reported increase in gamma frequency was not prominent in our V1-V4 site-pair selection). In principle, this was already achieved in **Fig. 2**, where we present the group-level repetition effects from 50 DCMs that could reproduce the data. With the sensitivity analysis in the two DCMs closest to the group-level repetition effect sizes, we aimed to understand whether all connections with significant group-level effects are meaningful and how they can bring about the empirical repetition-induced changes in gamma.

Informed by the PEB, we focused on those connections that showed significant group-level effects. Starting from the ‘late’ condition of the inverted DCMs, we examined the effects that the perturbation of individual connections in either a positive or negative direction can have on the DCM power and coherence. As in the data, each perturbation changed power and coherence spectra in the same direction (and in a qualitatively consistent way between the two monkeys, except in the case of the V1 ***s*** self-inhibition). This simplified our search for connection effects that were congruent with both the PEB and sensitivity analyses. We considered an effect congruent if (1) the effect was significant on the group level, (2) the perturbation of the respective connection produced changes in the gamma spectra of both representative DCMs, and (3) the sign of the group-level effect was the same as the direction of the perturbation that produced an increase in gamma power and coherence. The repetition effects that satisfied these conditions were: (1) the increase of V1 ***i***-to-***s*** and ***i***-to-***g*** inhibition, which were also the largest repetition effects in the PEB (47% and 30.6% respectively), (2) the increase of V1-to-V4 forward excitation, (3) the decrease of V4 ***g*** self-inhibition, (4) the increase of V4 ***g***-to-***s*** excitation, and (5) the increase of V4 ***i*** self-inhibition.

The sensitivity analysis was by design incomplete, in the sense that we did not consider all 50 DCMs or all parameter-change combinations, and somewhat counter to the very nature of a DCM, as we looked at the changes induced by individual connection strength modulations, whereas the DCMs and PEB integrated the repetition effects on all parameters simultaneously (taking into account their covariance). These shortcomings were inevitable since we wanted to zoom-in on specific models and their parameter constellations. The collective result of multiple connection modulations might look different than the mere sum of individual connection modulations. This complexity might be responsible, at least partly, for any inconsistencies between the two monkeys and for the inconsistencies between the PEB and the sensitivity analysis results. For example, only some of the connection strength modulations produced spectral changes that were in the direction expected by the PEB results. This does not mean that they were the only ones that should be considered or that they did not work together with the rest of the effects found in the PEB to reproduce the empirical gamma increases. Another example is that ***d*** self-inhibition was significant in the PEB but did not seem to affect gamma power or coherence in the sensitivity analysis. This lack of gamma changes with the perturbation of the efferents of ***d*** was expected from the design of the microcircuit, since the deep excitatory population and its associated parameters were included in the model to specifically capture lower frequency dynamics (Bastos et al., 2015a; Bastos et al., 2012). This decision was partly inspired by the finding that deep layers show less gamma and more alpha/beta (Buffalo et al., 2011) and that inter-areal directed influences in gamma are stronger in the bottom-up direction, whereas in beta they are stronger in the top-down direction (Bastos et al., 2015b; Michalareas et al., 2016; van Kerkoerle et al., 2014).

### 4.3 Stimulus repetition effects for few versus many repetitions

Two previous studies have investigated visually induced gamma for many stimulus repetitions and have reported strong gamma for the first presentation, rapidly decreasing gamma for the initial few (5-10) repetitions, and slowly increasing gamma for further repetitions (Peter et al., 2021; Stauch et al., 2021). The changes induced by the first presentation or the first few repetitions were not observed in the original study of Brunet et al. (2014) and in the task with gratings in Peter et al. (2021). What both of these latter experiments had in common was the presentation of grating stimuli to macaque monkeys that were highly overtrained on gratings. By contrast, in the task with natural images in Peter et al. (2021) and in the experiment of Stauch et al. (2021), the subjects had seen the stimuli a few times or for the first time respectively. Thus, it is possible that changes induced by the first presentation, or the first few repetitions vanish after extensive exposure to specific stimuli, and these changes might reflect effects of arousal. Arousal can be assessed through measurements of the pupil. Such measurements were performed in Stauch et al. (2021), and the observed pupil responses and gamma showed very similar dynamics across the first few stimulus presentations, suggesting that the strong gamma for the first presentation was related to arousal in response to a novel stimulus, and the decreasing gamma for the initial few presentations was related to rapidly decreasing arousal as those novel stimuli became more familiar. This is also consistent with the observation that the first presentation of novel natural stimuli to macaques often triggered saccades to those stimuli, suggesting image-triggered interest and potentially arousal, and that this effect rapidly diminished over the course of few repetitions (Peter et al., 2021). For the present study, we have used the dataset of Brunet et al. (2014). Therefore, our results are not concerned with those changes observed during the first presentation or the first few repetitions, but they are exclusively concerned with the increase in gamma induced when there are more than ten repetitions.

### 4.4 Stimulus predictability increases bottom-up precision and gamma

In the following paragraphs, we discuss repetition-induced changes and our findings in the context of the predictive coding framework. According to the predictive coding framework, the brain constitutes a generative model, under which sensory inputs are generated by real-world causes. This internal model is acquired through learning, is therefore continuously refined, and is used to infer the causes of sensory inputs from the inputs themselves. The difference between the actual sensory inputs and those predicted by the internal model gives rise to prediction errors. Prediction errors are propagated from lower to higher cortical areas by means of forward projections, i.e., they constitute the bottom-up inputs. Conversely, top-down inputs are predictions that are formed in higher areas and are projected back to lower areas to be subtracted from the local bottom-up inputs.

In predictive coding, prediction errors are weighted by their precision, a quantity that is equal to the inverse of their variance (Brown and Friston, 2012; Feldman and Friston, 2010; Friston and Kiebel, 2009). Inputs with high precision are inputs that have low variability, i.e., are more predictable. Predictability is linked to stimulus repetition. When a stimulus keeps repeating, its relative probability increases, i.e., its recurrence becomes increasingly predictable. This means that over the course of many repetitions, the predictability of bottom-up stimuli, and the precision of associated prediction errors, should increase monotonically until it plateaus. As described in the previous section, this is exactly what has been reported for gamma power as a function of many repetitions in all previous experiments that report such changes in gamma (Brunet et al., 2014; Peter et al., 2021; Stauch et al., 2021). Therefore, gamma oscillations might mediate increases in predictability or precision by modulating the gain of bottom-up prediction errors. Indeed, gamma synchronization has been related to an increased precision in neuronal signals; namely, a decrease in noise correlations to near-zero values in macaque V1 (Womelsdorf et al., 2012). Furthermore, neuronal synchronization has been proposed to increase the postsynaptic impact or gain of spikes on a receiving population (Bazhenov et al., 2005; Gotts, 2003; Gotts et al., 2012). Specifically, inter-areal neuronal synchronization in the gamma-frequency range has been related to the feedforward communication of bottom-up inputs (Bastos et al., 2015a; Bastos et al., 2015b; Bastos et al., 2012; Fries, 2015; Friston et al., 2015; Michalareas et al., 2016; van Kerkoerle et al., 2014; Vezoli et al., 2021).

Another parameter that determines the precision of bottom-up input is the spatial predictability of the respective sensory stimuli. In stimuli that are spatially uniform, in their color or structure, one part can be predicted from other parts. Intriguingly, the spatial predictability of visual stimuli is strongly related to the strength of the induced gamma in macaque V1. Gamma induced by a patch of grating or of uniform color in the RF is enhanced if the same grating or color is extended into the RF surround (Gieselmann and Thiele, 2022, 2008; Jia and Kohn, 2011)(Peter et al., 2019). Similarly, gamma induced by a patch of a natural image is enhanced when the same stimulus is extended into the RF surround and when the part inside the RF is spatially predictable from the part outside the RF (Uran et al., 2022).

The role of precision in weighting prediction errors within the predictive coding framework leads to a second prediction. That is, precision should modulate the gain of the populations that encode and transmit prediction errors to higher areas. Indeed, this is the effect that the DCM revealed: (1) a decrease in the self-inhibition of the V1 ***s*** population, which projects prediction errors to higher areas, and (2) a decrease in the self-inhibition of the V4 ***g*** population, which receives the prediction errors from V1 and relays them to other V4 populations. In the model, the self-inhibitory connections control the gain of the respective populations, and a decrease in self-inhibition means an increase in the gain of the population. Moreover, as mentioned above, the self-inhibition decrease in both populations can result in an increase in local gamma power and inter-areal gamma coherence as shown in the sensitivity of representative DCMs from each monkey. Thus, these two effects that we report here are in line with the proposed role of precision and, more specifically gamma oscillations, within the predictive coding framework.

A similar effect of predictability has been reported before in a DCM study (Auksztulewicz et al., 2017) of MEG responses in the auditory cortex to repeated sequence of tones (Barascud et al., 2016). Two factors that increase stimulus predictability, i.e., the repetition of regular versus random sequences and a smaller versus larger number of possible tones that could be used to form a sequence, had a positive change on MEG responses between 8 and 128 Hz. This positive change in the responses was modelled by an increase in the gain of pyramidal cells in the auditory cortex and in the inferior temporal gyrus (Auksztulewicz et al., 2017).

In addition to the aforementioned self-inhibition decreases, we observed an increase in inhibitory connection strengths within the V1 microcircuit, i.e., from the ***i*** to the ***g*** and ***s*** populations, from the ***s*** to the ***g*** population and in the self-inhibition of ***g***. The strength of inhibitory connections might be increased by an increase in the gamma-band synchronization of the respective neurons, and this has indeed been reported as an effect of stimulus repetition for putative inhibitory (narrow-spiking) cells in V4 (Brunet et al., 2014). An increase in inhibitory connection strengths with repetition was also proposed later by Peter et al. (2021) and Stauch et al. (2021): it would need to target specifically the more weakly-driven principal cells, eventually inhibiting them from firing within a gamma cycle. Our mesoscopic-level model is not able to differentiate between strongly and weakly driven excitatory populations. Therefore, an obvious extension of the current model will include both strongly and weakly driven excitatory populations. Ideally, strongly and weakly driven excitatory neurons would then also be experimentally identified and recorded. The same holds for their laminar position, which would allow to use the laminar experimental data to explicitly constrain the model populations of the corresponding layers.

## Acknowledgements

The authors would like to thank Eleni Psarou for the invaluable scientific input during all stages of this project and Peter Zeidman for discussions and advice on DCM methods.

CAB was supported by the FLAG-ERA JTC-2023 project MONAD (co-financed by ZonMw, project number JPND-10960352310003). KJF was supported by funding for the Wellcome Centre for Human Neuroimaging (Ref: 205103/Z/16/Z). PF was supported by DFG (FR2557/2-1, FR2557/5-1, FR2557/ 7-1), EU (HEALTH-F2-2008-200728-BrainSynch, FP7-604102-HBP), a European Young Investigator Award, National Institutes of Health (1U54MH091657-WU-Minn-Consortium-HCP).

## CRediT author statement

CK: conceptualization, methodology, software, validation, formal analysis, writing - original draft, writing - review and editing, visualization. CAB: data curation, investigation, writing - review and editing. KJF: conceptualization, methodology, writing - original draft, writing - review and editing, supervision. PF: conceptualization, investigation, resources, writing - original draft, writing - review and editing, supervision, project administration, funding acquisition.

---

**Figure S1.**
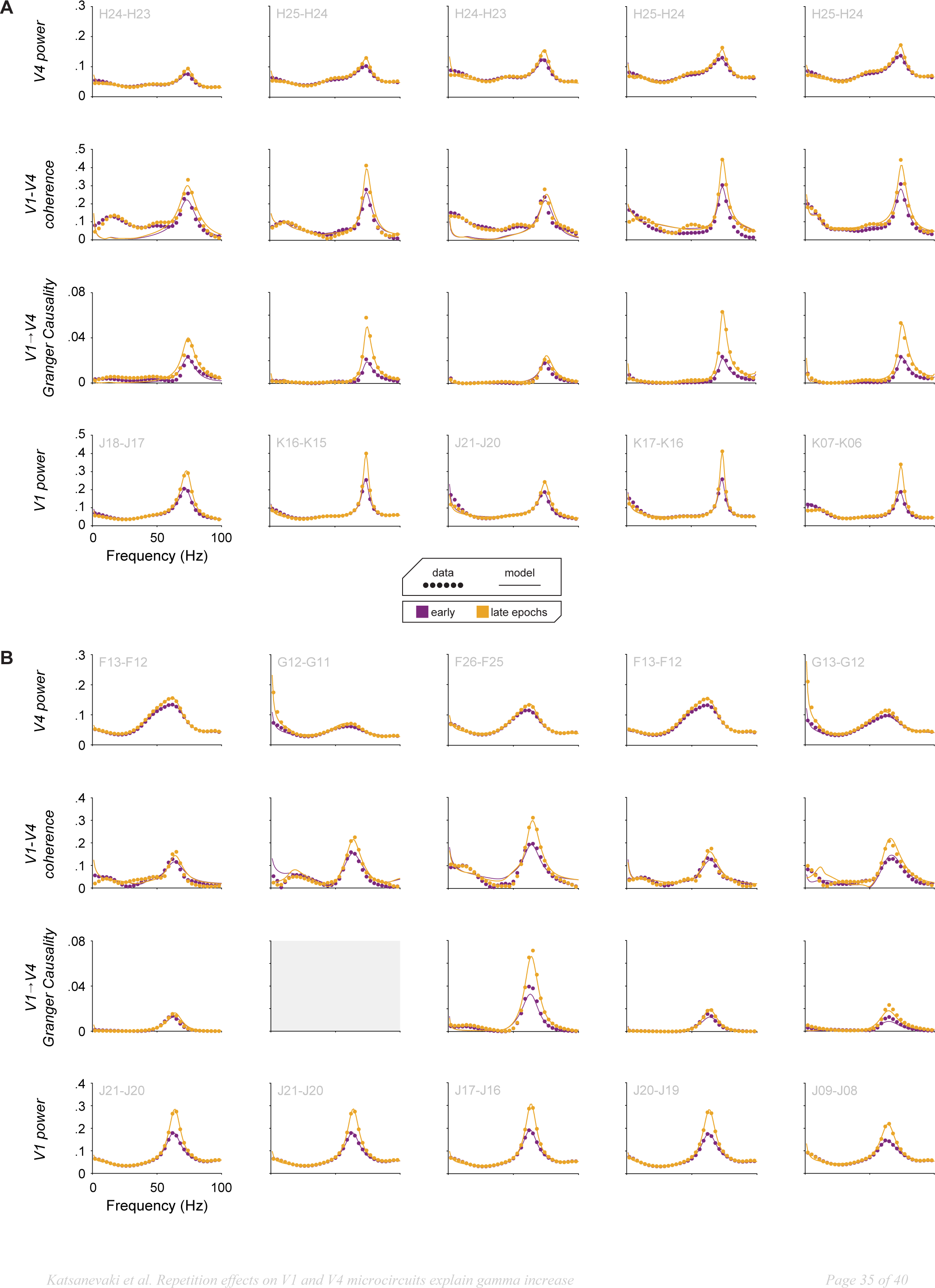
**(A)** Empirical (*circles*) and predicted (*lines*) spectra of V1 power (*bottom*), V1-to-V4 GC, V1-V4 coherence and V4 power (*top*) of 5 randomly selected V1-V4 pairs of monkey K. Grey text indicates the bipolar site label to show sites that are presented multiple times. **(B)** Same as above, but for 5 randomly selected pairs from monkey P. For one pair from monkey P, the GC calculation did not converge and is therefore omitted.

**Figure S2.**
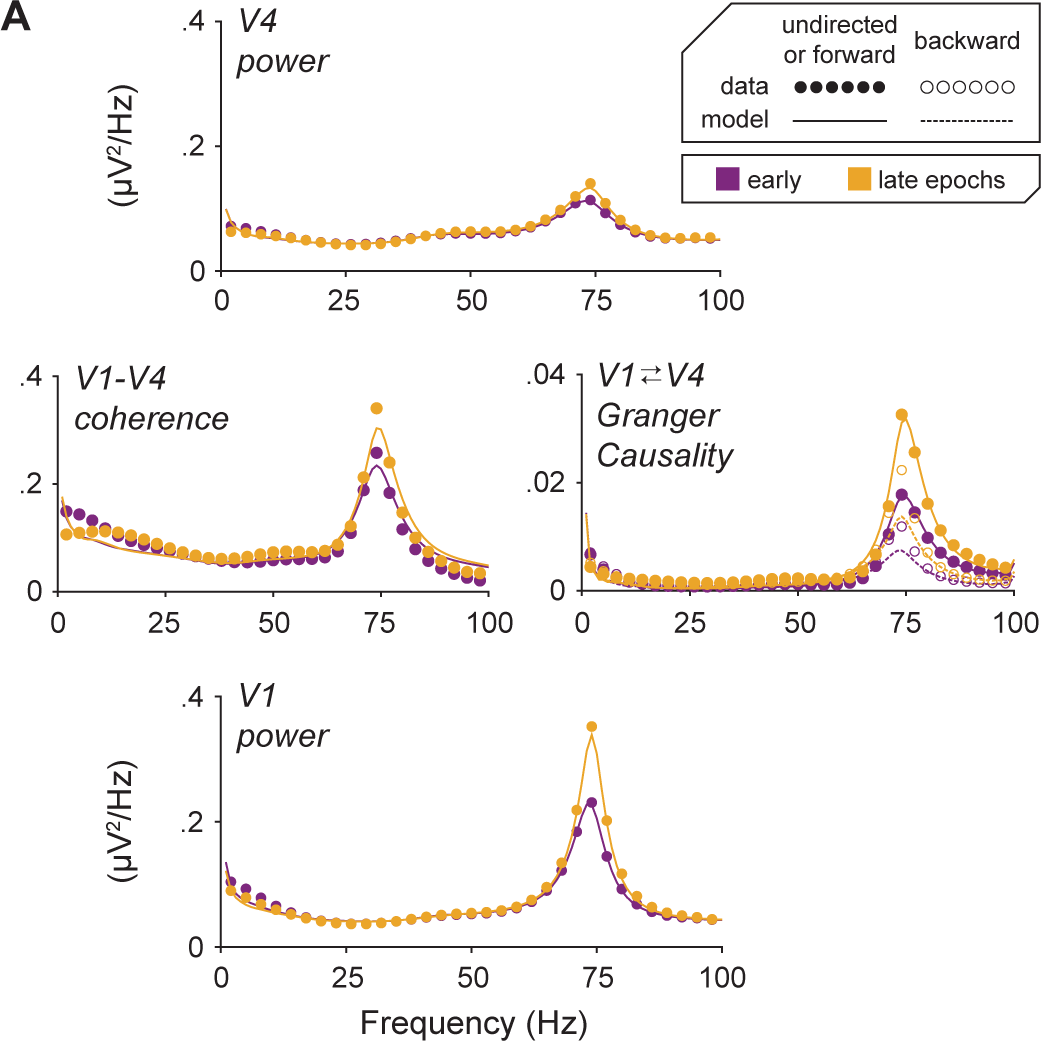
Empirical (*circles*) and predicted (*lines*) spectra of V1 power (*bottom*), V1-V4 coherence and GC (*middle*) and V4 power (*top*) averaged over all V1-V4 site pairs and DCMs of monkey K. Conventions as in Fig. 2A.

**Figure S3.**
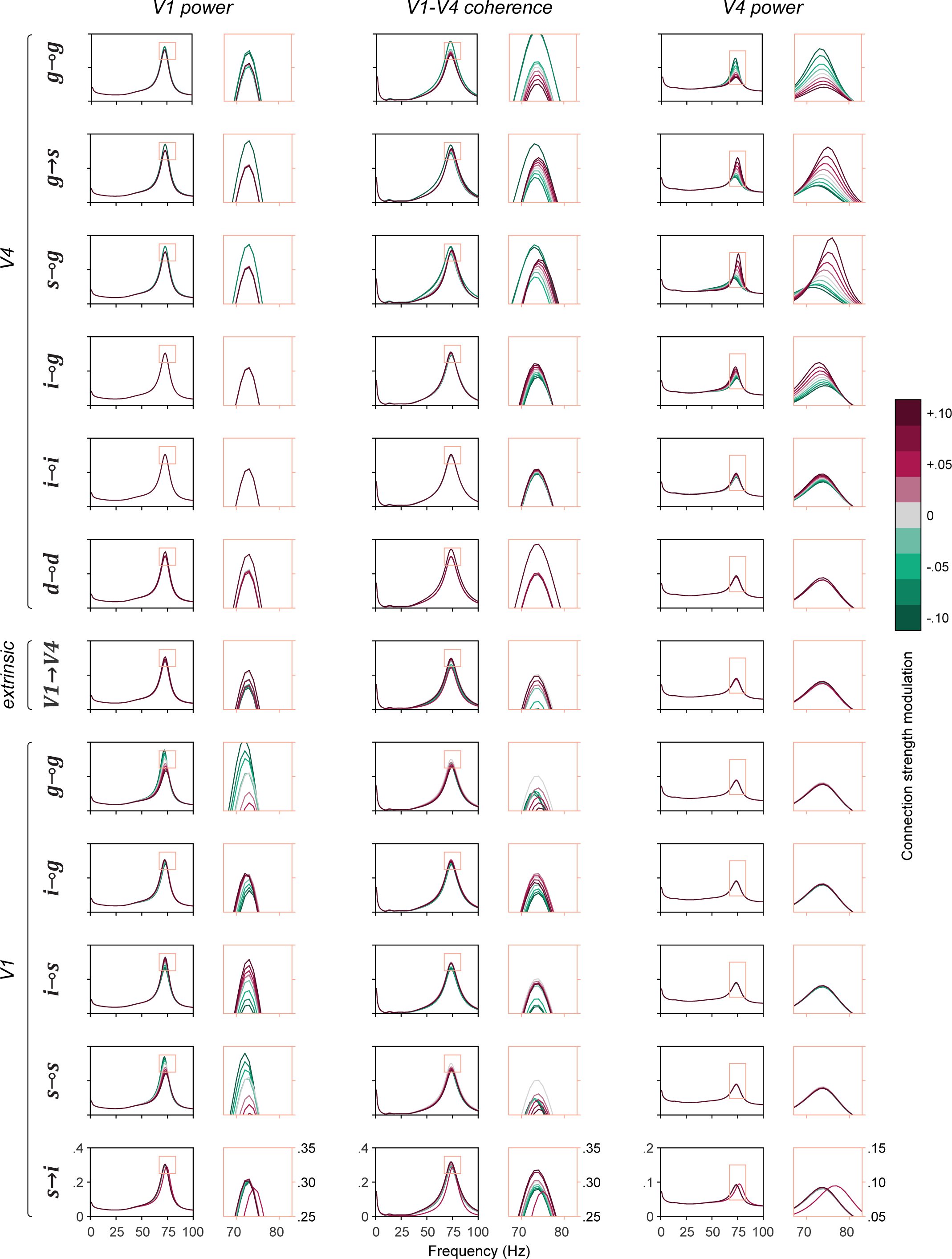
Sensitivity of V1 power (*left*), coherence (*middle*) and V4 power (*right*) spectra to small perturbations of the strength of individual connections in a representative DCM of monkey K. Conventions as in Fig. 3.

